# Ancient DNA from shells reveals delayed genomic erosion and rapid immune adaptation in the critically endangered black abalone

**DOI:** 10.64898/2025.12.01.690857

**Authors:** T. Brock Wooldridge, Joshua D. Kapp, Sarah M. Ford, William E. Seligmann, Holland C. Conwell, Talia Tzadikario, Jonas Oppenheimer, Zachary G. Anderson, Alan Le Moan, Alicia Abadía-Cardoso, Pete Raimondi, Beth A. Shapiro

## Abstract

Predicting the genetic consequences of population decline is a major problem in conservation genomics. Time lags following demographic bottlenecks can delay genomic erosion and make it difficult to determine a population’s current and future risk, especially when pre-bottleneck genomic baselines are unavailable. Black abalone (*Haliotis cracherodii*) suffered a severe disease bottleneck in the 1980s, resulting in an estimated 99% population decline. However, recent work found surprisingly high genetic diversity and little population structure in current black abalone populations, raising questions of whether genomic erosion has been delayed. To investigate this, we applied ancient DNA methods to pre-bottleneck abalone shells, generating 59 whole genomes including one 34-fold coverage genome from a 1,500-year-old specimen. These data show that heterozygosity, runs of homozygosity, genetic load and population structure remained stable up to and following the bottleneck. Simulations reveal that this stability is consistent with even severe bottleneck scenarios because too few generations have lapsed since the decline. Projections suggest that future genomic erosion may be avoided even in limited recovery scenarios. Following the bottleneck we observe widespread balancing selection at genes with immune function, along with parallel increases of two inversions on separate chromosomes that are in linkage disequilibrium, where the disease bottleneck was most severe. Altogether, these findings explain why genomic change has thus far been limited, outline recovery scenarios that minimize genomic erosion, and identify loci likely that may harbor adaptive variation key to the success of future black abalone populations.

## Introduction

Rapidly declining populations are susceptible to genomic erosion, or the combined effects of reduced diversity, inbreeding, and mounting genetic load that can increase extinction risk ^1^. However, the timing and magnitude of these effects is far from predictable, as illustrated by the recent history of the black abalone *Haliotis cracherodii*. This intertidal mollusk was once abundant along the west coast of North America, serving as a common food item and cultural keystone species ^2^. In 1985, a bacterial disease known as “Withering Syndrome” (WS) first appeared and drove an estimated ∼99% decline within just a few years ^3,4^. Despite this well-documented collapse, the genomic consequences of this rapid decline are not clear ^5–7^. Black abalone across California have a high effective population size (Ne ∼ 300,000; ^8^, show extraordinarily high contemporary genetic diversity, and exhibit no evidence for genetic isolation between sites ^7^. All of these features appear inconsistent with expectations for a population that recently underwent near-extinction event ^9–11^. Without a historical baseline it is difficult to ascertain if any current genomic patterns have been affected by the recent bottleneck, which can impact how management strategies proceed.

It is common to see a mismatch between census population declines and population genomic data. Both empirical and theoretical work shows that many of the signatures of genomic erosion may be slow to reflect population decline ^12,13^. This delay is often referred to as a “time lag” ^14^ or “extinction debt” ^15^.

Compounding this, reconstructing recent changes based solely on modern genomes is challenging ^16^. Temporal genomics, or the time series analysis of genomes ^17,18^, offers a powerful way to directly measure genomic erosion over time ^19,20^. Temporal genomic studies of bottlenecks have revealed both expected ^21–25^ and unexpected patterns, including limited genomic erosion ^13,17,26^ or increases in diversity ^27^. Emerging data and theory suggest that time lags are linked to life history and long-term demographic processes ^28^. The life history traits of black abalone make a time lag probable - they can live up to 30 years, reach sexual maturity at 4 years, and have highly overlapping generations ^28–30^. However, a predictive framework that could estimate the time lag effect remains elusive. Without this framework, pre-bottleneck genomes provide the best way to compare current populations to a pre-collapse baseline and forecast future change .

Temporal genomics also provides the opportunity to track allele frequency changes through time, which may indicate natural selection. Temporal genomics studies have revealed genes underpinning adaptation to rapid selection events in wild populations, including cold tolerance in *Anolis* lizards over a single generation ^31^, and viral resistance in rabbits over tens of generations ^32^ . In other cases, signals of adaptation have been difficult to detect, which might be attributed to subtle signals of polygenic adaptation or the lack of heritable adaptive variation ^17^. In black abalone there is some evidence for heritable differences in Withering Syndrome susceptibility ^33^ , which is further supported by complementary experiments in other abalone species ^34,35^. If WS resistance has indeed evolved, then it’s possible that pre-and post-bottleneck genomic comparisons could reveal the gene(s) underlying WS adaptation. Previous work did not identify any obvious targets of selection aside from a chromosomal inversion associated with latitude ^7^. This inversion remains a compelling candidate as latitude is associated with both temperature and WS spread ^3,36^, and chromosomal inversions are frequently implicated in local adaptation ^37^. The identification of any loci showing local adaptation over space or time will be key in guiding the translocation plans that are only just beginning ^38^.

Genomes from pre-bottleneck black abalone are needed to measure genomic erosion and selection, but generating genomic data from ancient and historic mollusk specimens is challenging. DNA damage can accumulate quickly through environmental and chemical mechanisms if preservation (e.g. freezing) is insufficient to slow down this process ^39^. Because shells and dry preparations account for 91% of all abalone in Malacology (mollusk) collections ^40^, specimens available for genome sequencing are limited to sample types with inherently damaged DNA. Specialized sample processing and analysis methods are needed to handle the small quantity of degraded DNA present in these specimens. Thus far, less than 100 mollusk shells across four studies ^41–44^ have been processed for whole genome DNA analysis (i.e. shotgun sequencing), none of which have generated multi-fold nuclear genome coverage. In contrast, for vertebrate species it is common to generate whole genomes for hundreds of ancient individuals in a single study ^45,46^. In mollusks, DNA is entrapped during the shell formation process and higher organic content in the shell structure underlies greater DNA preservation ^47,48^. Consistent with this, endogenous DNA content has been reported to be as high as 30% for some specimens ^44^. Given this evidence and a growing understanding of how different shell features preserve DNA ^44^, it seems feasible to conduct population genomic studies based on large sets of shells.

Here, we successfully apply best practices in ancient DNA sequencing to a time series of shells spanning 1500 years of black abalone history. Using multi-fold coverage shell genomes, we are able to directly assess changes in genetic diversity, inbreeding, population structure, and natural selection over time, particularly in relation to the Withering Syndrome bottleneck. We then use these findings to predict scenarios of future genomic erosion and identify variation that could be adaptive. These findings are key to managing the recovery of this critically endangered species.

## Results

### Temporal genomics of black abalone

We sequenced whole genomes from 59 black abalone shells spanning the species’ range and the past 1500 years (Fig. 1). Most shell specimens are dated from 1914-1979, prior to the first mass mortalities due to Withering Syndrome in 1986 ^4^. We also included one shell from the present day to serve as a control sample. To obtain these data, we took ancient DNA protocols optimized for bone and applied them to abalone shell fragments ^49,50^. Specifically, we pulverized a ∼50mg piece of each shell, pre-treated the resulting powder with bleach ^44^, extracted DNA from the powder using a silica spin column for small DNA fragment recovery ^50^, and generated single-stranded DNA libraries from each extraction ^49^. All steps were performed in a dedicated ancient DNA lab to minimize contamination. Sequencing libraries resulting from this pipeline ranged in endogenous DNA content from 0.5% to 68.6%, although the median content amongst 20th century museum shells was 49.5%. DNA damage also scaled with sample age. For the samples dated 1500 BP, cytosine deamination frequency at the 5’ termini and average mapped fragment length were 21% and 60 bp, respectively. These values were 4.2% and 104 bp in the 20th century shells, indicating far lower rates of damage (Fig. S1). Our final shell dataset consisted of 44 samples sequenced at ∼2X coverage and 15 sequenced at 20X or greater coverage. This latter group included a 34X genome from a shell midden dated to 1500 BP.

**Figure 1.**
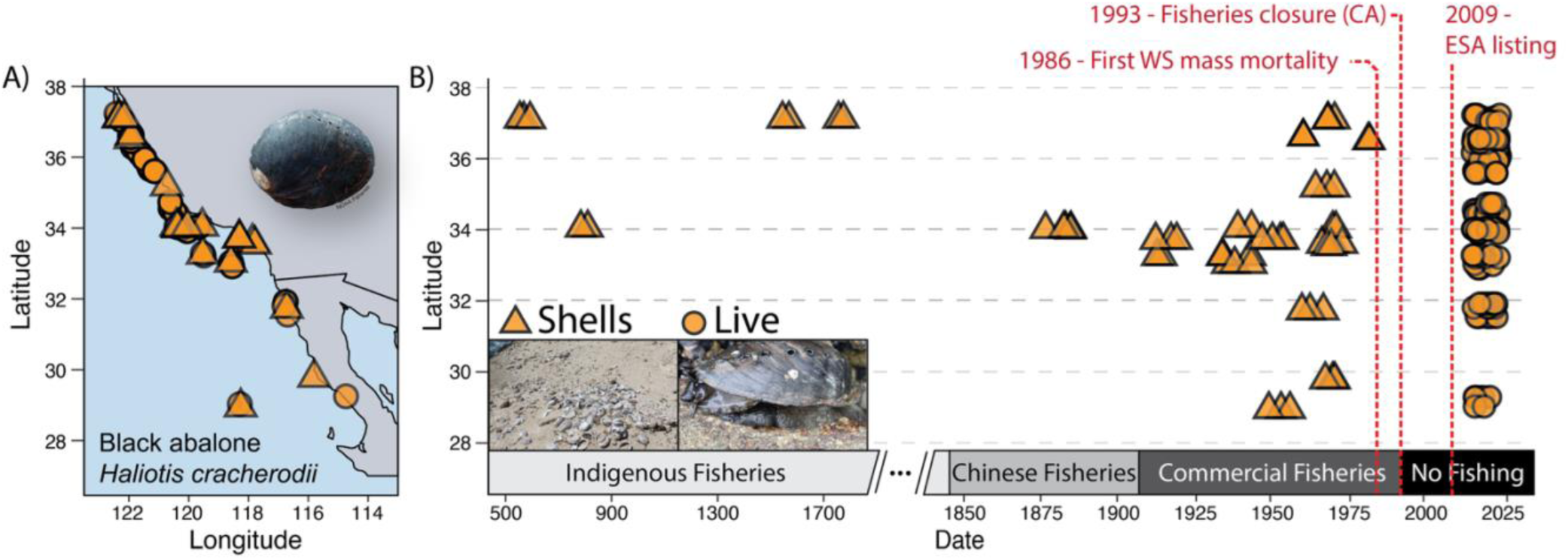
**Study design**. A) Distribution of shells (triangles) and live specimens (circles) sequenced in this study. B) Median estimated ages of sequenced shells. Y axis position corresponds to latitude in panel A, and bands on the x-axis indicate the active fisheries when these specimens were alive.

Finally, we combined these ancient and historic shell genomes with 138 modern black abalone genomes published in Wooldridge et al. (2024) and 16 modern genomes from the Baja California range, new to this study.

### No change in genetic diversity, inbreeding, or load following the Withering Syndrome bottleneck

A direct comparison of historic and modern black abalone shows no change in genetic diversity over time (Fig. 2). We found that individual heterozygosity remained consistent at 0.45-0.50% across all time periods (Fig. 2A). Similarly, the fraction of the genome in runs of homozygosity (FROH) remained below 1% across all time periods and was 0% for the majority of individuals, although the ‘Commercial Fisheries’ periods exhibited substantially more variation than all other periods (Fig. 2B). Finally, we also recorded limited change in either masked (Fig. 2C) or realized (Fig. 2D) genetic load across time periods ^51^. Overall, we see no substantial effect of the Withering Syndrome bottleneck on any of these measures of genetic diversity or inbreeding depression.

**Figure 2.**
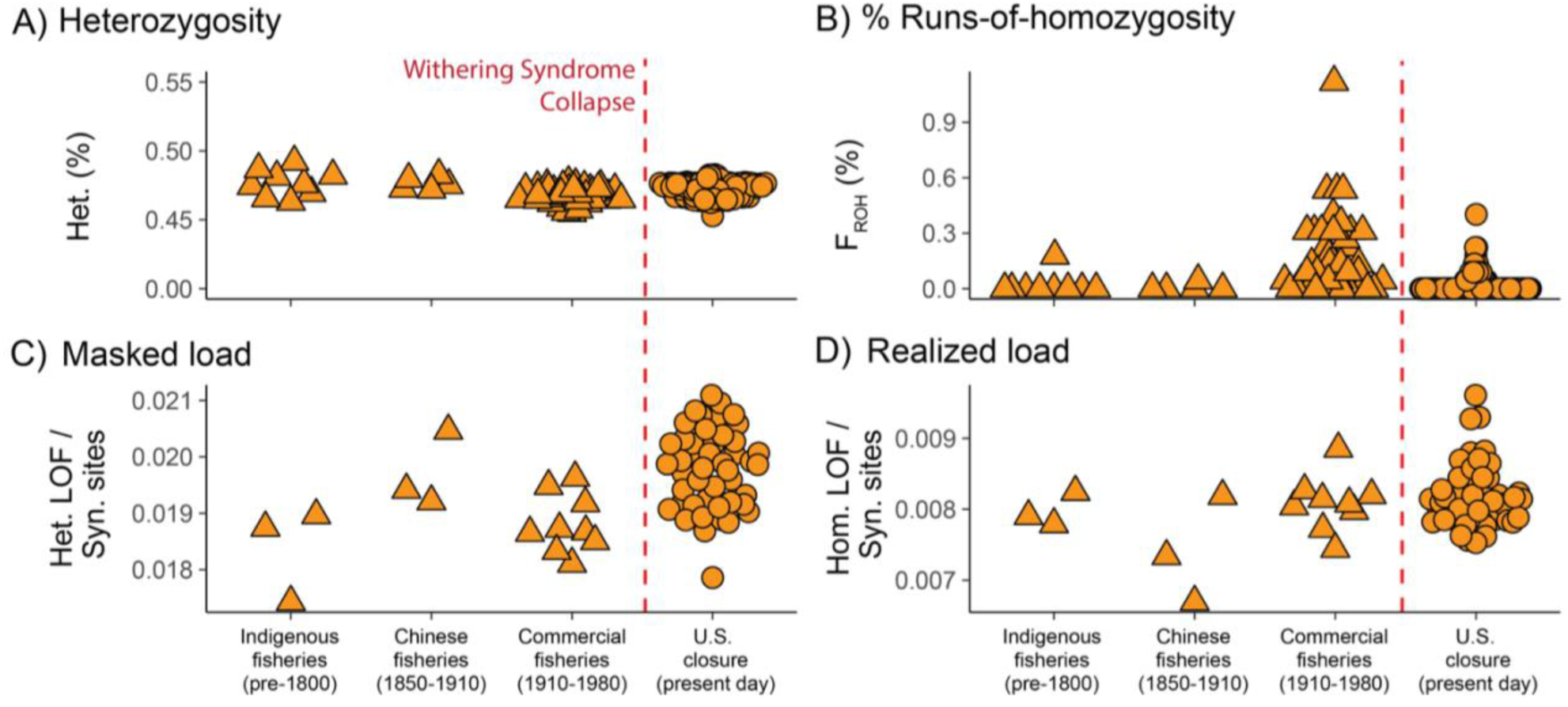
**Metrics of genetic diversity and inbreeding depression over time**. Both heterozygosity (**A**) and FROH (**B**) are calculated to account for sequencing coverage and DNA damage ^52^. Masked load (**C**) and realized load (**D**) are calculated only on high coverage (>15X) individuals using transversion polymorphisms.

### Subtle changes in population structure occur following Withering Syndrome Bottleneck

We observe only subtle changes in population structure between pre- and post-bottleneck black abalone. An initial genetic PCA showed that all samples fell into one of three discrete clusters with some notable outliers (Fig. 3A). Upon closer inspection, we found that all outliers belonged to one of the following groups: 1) shells from a ∼1500 BP site at the northern end of the range (blue circle, Fig. 3A), 2) historic and modern samples from Isla Guadalupe where a putative subspecies has been described ^53^ (red circle, Fig. 3A), 3) historic and modern samples from Faro San Jose, representing the southern end of the species’ continental range (purple circle, Fig. 3A). Samples from this last group were outliers along PC2, but along PC1 aligned with the three primary clusters. Within the three dense clusters along PC1 we see no association with geography or sample age that is not better explained by sequencing depth.

**Figure 3.**
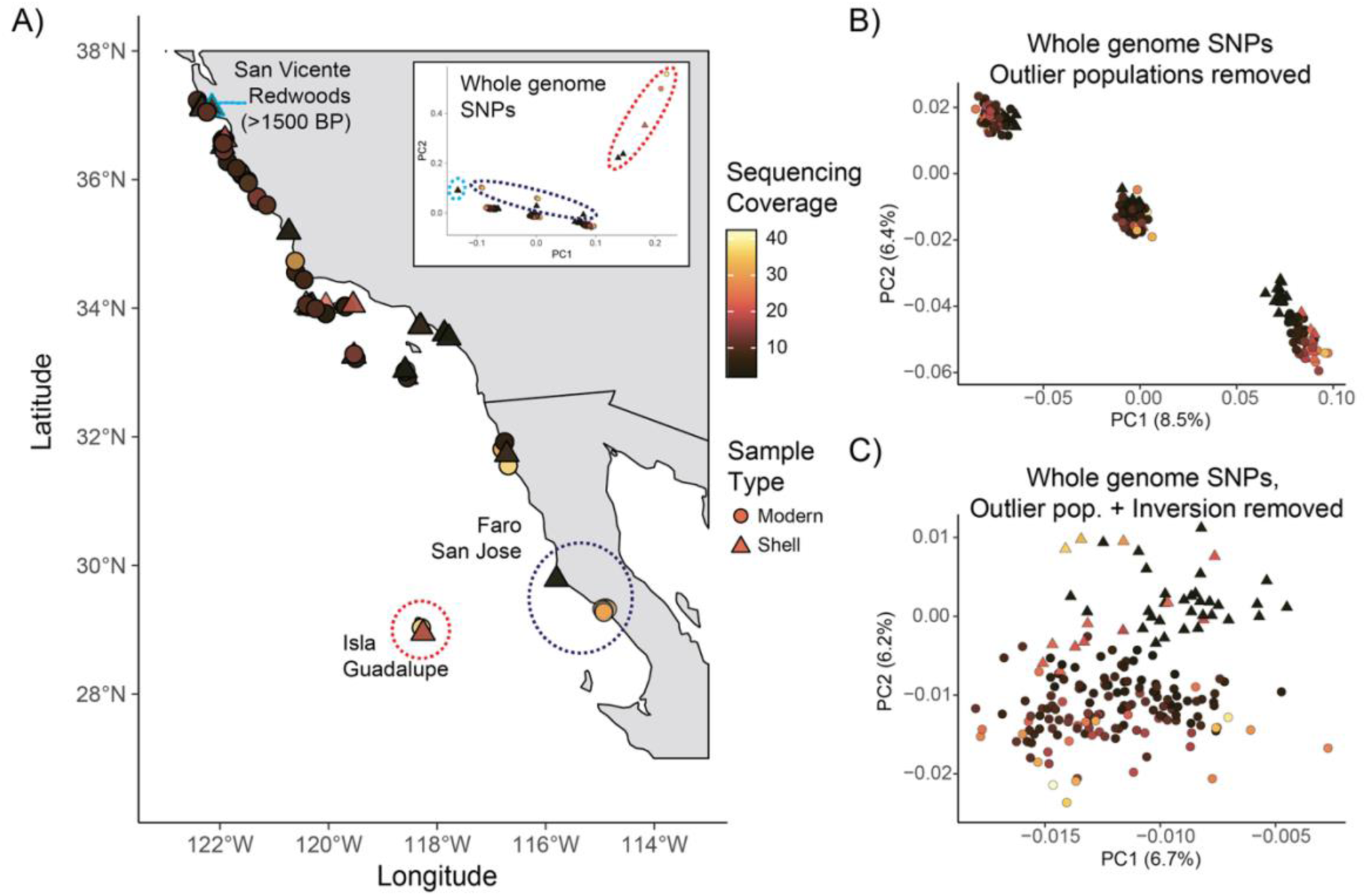
**Population structure over time**. **A)** Sample map, inset shows whole-genome PCA. Groupings indicated by dotted lines show how outliers correspond to geographic or temporal outliers. **B)** Same PCA as in panel A), but with outlier populations removed. **C)** PCA excluding outliers populations and SNPs within the 31 Mb chr4 inversion described by Wooldridge et al. (2024).

The three PC1 clusters recapitulate the structure described in Wooldridge et al. (2024) that was attributed to a 31 Mb chromosomal inversion on scaffold 4 (Fig. 3B). After removing SNPs within the boundaries of the inversion locus, the three clusters disappear and only a diffuse cloud of samples remains (Fig. 3C). Within this cloud we see marginal separation of historic from modern samples along PC2, and this separation is not driven by sequencing coverage. Consistent with this, median Hudson’s FST between pre-bottleneck (1914-1979) and post-bottleneck samples is only 0.008, and genetic diversity (π) stays nearly constant at 0.82% and 0.81%, respectively (Fig. S2). Taken together, these results indicate limited genetic divergence between pre- and post-bottleneck black abalone.

### A time-lag effect on genomic erosion

Simulations show that the stability in diversity, inbreeding, and structure we observe is possible even following a severe bottleneck (Fig. 4). To determine this, we first summarized the average change in heterozygosity, FROH and FST between pre-bottleneck, ‘Commercial Fisheries’ era samples (1914-1979) and post-bottleneck samples. We then simulated populations undergoing a bottleneck followed by migration between newly formed subpopulations (Fig. S3). The declines and recoveries we simulated were informed by records of Withering Syndrome impacts and recent ecological surveys (Fig. S4) ^4^. We see that our time-series measurements are close to simulated ones at 10 generations post-bottleneck, which is roughly the number of generations since WS first appeared (Fig. 4). This is true even if we simulate a population reduction of 99.9%, which is more extreme than most decline estimates ^4^ (Fig. 4B). Our simulations also show limited divergence between the newly formed subpopulations that arise post-bottleneck. These results indicate that a time-lag may explain why there is little evidence for genomic erosion in the present day.

**Figure 4.**
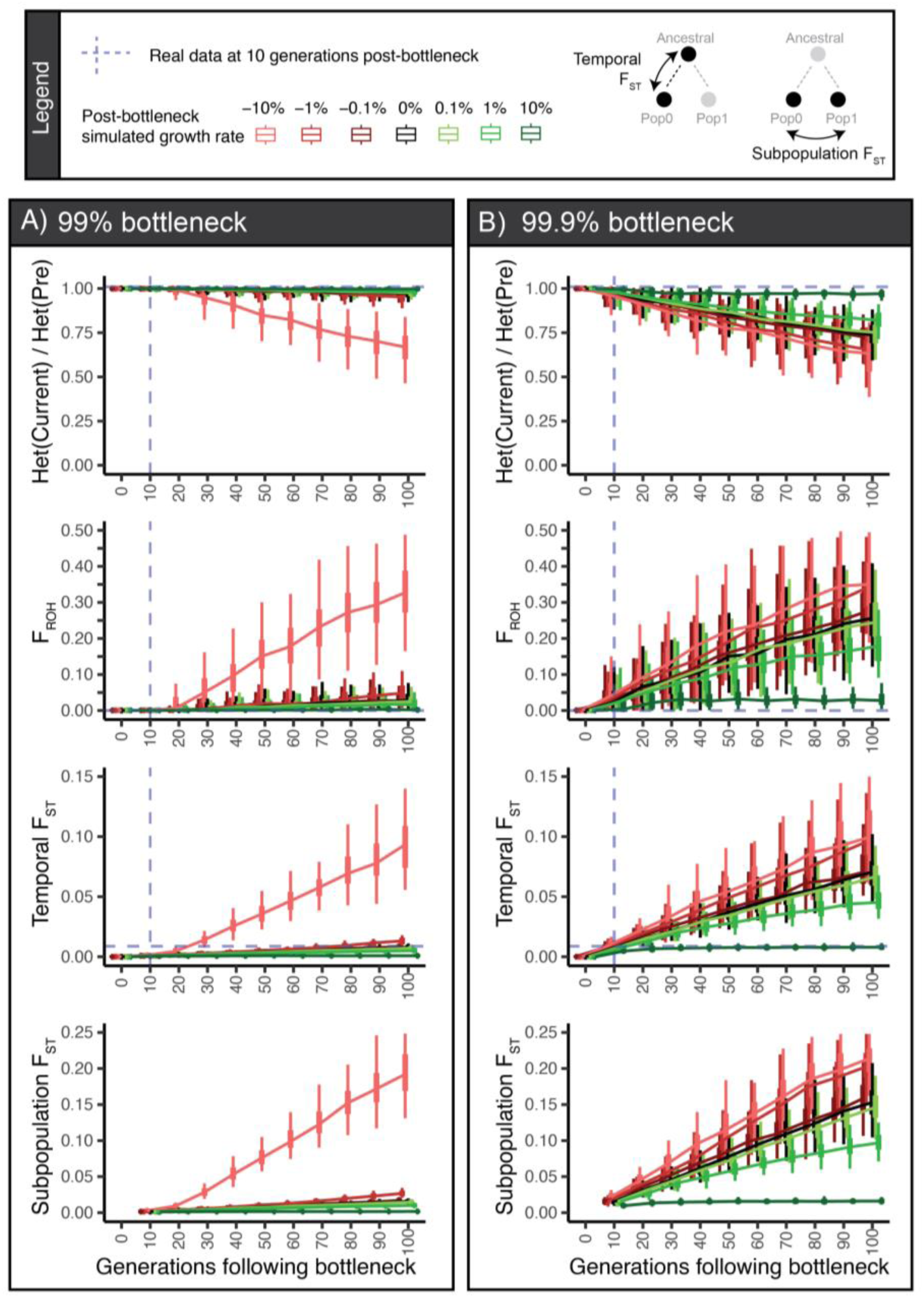
Simulated effects of a bottleneck and recovery on inbreeding metrics. We simulated two bottleneck intensities, **A)** 99% of former size, and **B)** 99.9% of the former size. All growth and decline ceases if Ne returns to the pre-bottleneck Ne (∼3.0e5) or drops to 100 individuals. See Fig. S3 for model and metric details.

However, as post-bottleneck time increases in our simulations, we do begin seeing clear signs of genomic erosion. This is most evident in the more extreme 99.9% bottleneck simulation (Fig. 4B). Only recovery rates of 1% or greater are able to stabilize heterozygosity, FROH, and both measures of FST in the long term (Fig. S5B). In contrast, the 99% bottleneck does create significant short term (Fig. 4A) or long term (Fig. S5A) genomic erosion even if population recovery is 0%. These projections highlight how much the initial severity of the bottleneck determines outcomes at short and long timescales. At 99.9% intensity, delayed genomic erosion is likely unless recovery is significant and sustained. A lesser 99% bottleneck, which reflects the range-wide summary of WS decline ^4^, is less likely to result in genomic erosion even if recovery is negligible.

### Evidence for recent selection at immune loci

We identified large genomic regions exhibiting post-bottleneck balancing selection (Fig. 5). For our selection analyses we removed the oldest samples, focusing just on pre-bottleneck (1914-1979) and post-bottleneck individuals, then calculated the difference in π between both time periods (‘πpost-pre’) along with Hudson’s FST in 50 kb windows. Windows in the top 1% of FST and πpost-pre correspond to genetic divergence between the time periods accompanied by an increase in genetic diversity towards the present day. We tentatively label these windows as under balancing selection ^54,55^, although we acknowledge that other selective processes could be responsible ^56^. We identify 50 outlier windows with these features clustered in large islands on scaffolds 7, 12, 14, and 19, containing an array of innate immunity and gamete recognition genes. These islands are significantly enriched for scavenger receptor genes in particular (p = 0; see Methods). One of these scavenger receptors, *DMBT1*, is known to be involved in immune responses to bacterial challenges in abalone ^57^. A phylogeny of the *DMBT1* outlier region confirms that pre-bottleneck diversity is a small subset of modern diversity at this locus (Fig. 5E) Windows in the top 1% of FST and the bottom 1% of πpost-pre may indicate a recent selective sweep, as these genomic regions have both diverged and decreased in relative diversity towards the present. In contrast to the analyses above, we identified only 11 windows exhibiting selective sweep characteristics. Of the 17 genes present within these windows, one gene - a homolog to *SVEP1* (Sushi, von Willebrand factor type A, EGF and pentraxin domain-containing protein 1) - is a compelling candidate. *SVEP1* has been associated with total body weight in a QTL study of South African abalone ^58^, and also acts as a common shell matrix protein and has immune function in mollusks ^59,60^. A phylogeny of the outlier region overlapping *SVEP1* shows three distinct clades of short branches, suggesting multiple selected haplotypes and a soft sweep (Fig. 5D). Although not an FST outlier, we also see a dramatic drop in diversity at the 31

**Figure 5.**
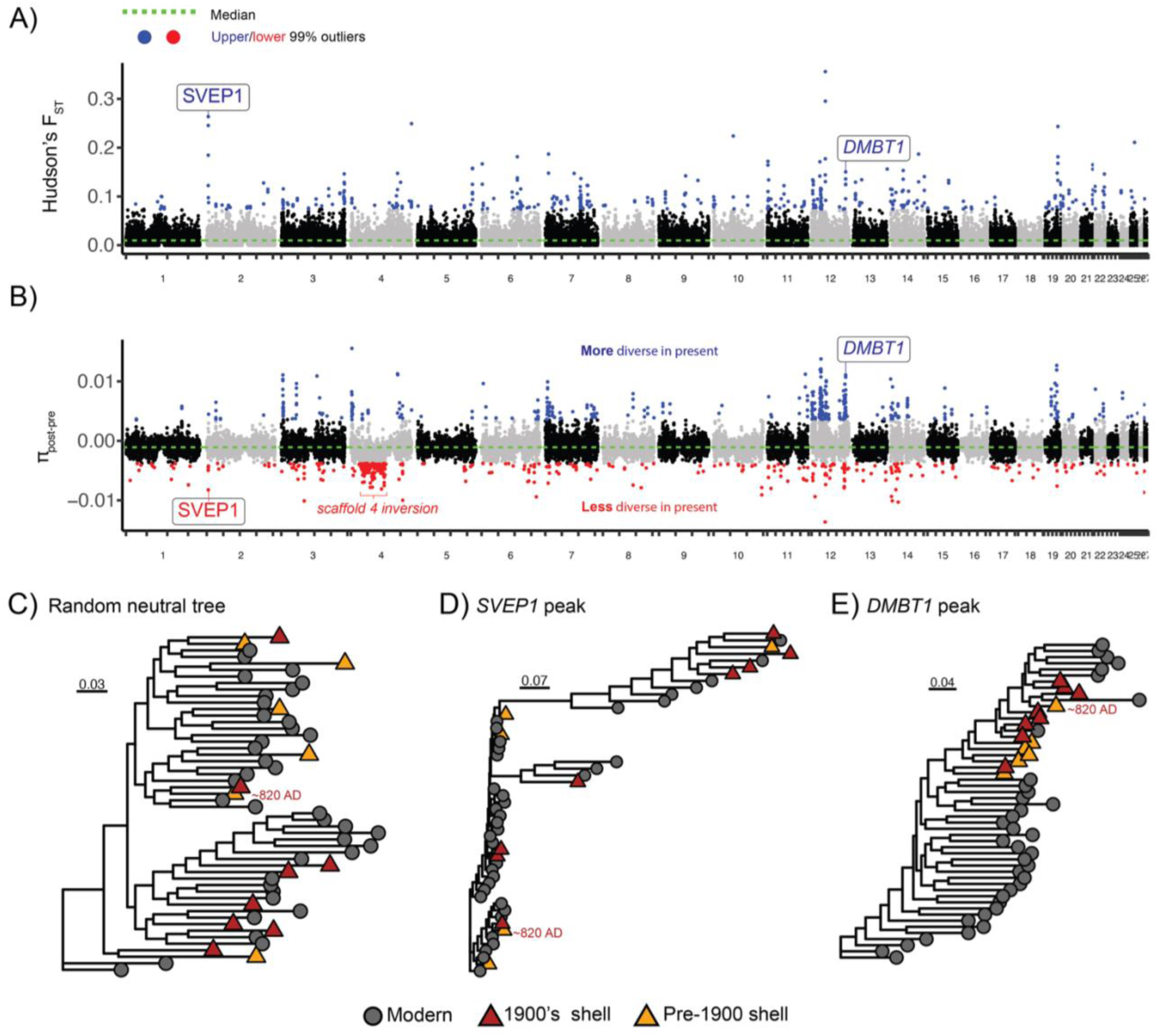
Evidence for natural selection over time. **A)** FST and **B)** πpost-pre between pre-bottleneck (1914-1979) and post-bottleneck samples. Each point in A) & B) represents a 50 kb x 25 kb sliding window. **C)-E)** Maximum likelihood phylogenies of 50 kb windows for a representative neutral region (C), the selective sweep at *SVEP1* (D) and balancing selection at *DMBT1* (E).

Mb inversion on scaffold 4. Both the inverted and reference alleles appear to have lost diversity after the bottleneck (Fig. S6). In sum, balancing selection in association with the Withering Syndrome bottleneck seems to have been more common than positive selection, although classic selective sweeps occurring on this timescale are difficult to detect (see Discussion). Nevertheless, these analyses uncovered several genes that are plausibly involved in black abalone immune adaptation over recent generations. We have included a list of these genes and the relevant literature in Extended Data Table 1.

### Parallel increases of two inversion alleles following the Withering Syndrome bottleneck

Two inversions on separate chromosomes showed similar frequency shifts following the WS bottleneck (Fig. 6). Initial genotyping of the two inversions, one on scaffold 4 and the other scaffold 9 (Fig. S8), suggested that both were more common at higher latitudes. To formally test this, we fitted stable, linear, and sigmoid clines to inversion presence or absence across latitude. Model comparisons showed that sigmoid clines were the best fit for each inversion in both pre- and post-bottleneck time periods, although stable clines also had high support for the scaffold 9 inversion (Table S1). The inflection point of each cline remained at or just north of Pt. Conception, a common biogeographic barrier in marine systems ^61^ (Fig 6A-B; Table S1). While a clinal pattern is always present, south of Pt. Conception both inversions doubled in frequency following the bottleneck (Fig. 6A-B & Fig. S7). Range-wide post-bottleneck increases of the scaffold 4 and scaffold 9 inversions amount to 23.9% and 17.6%; these values fall in the top 1.5% of genome-wide allele frequency shifts. In addition to overall changes in frequency, polymorphism within both the inverted and collinear alleles drops following the bottleneck (Fig. S6).

**Figure 6.**
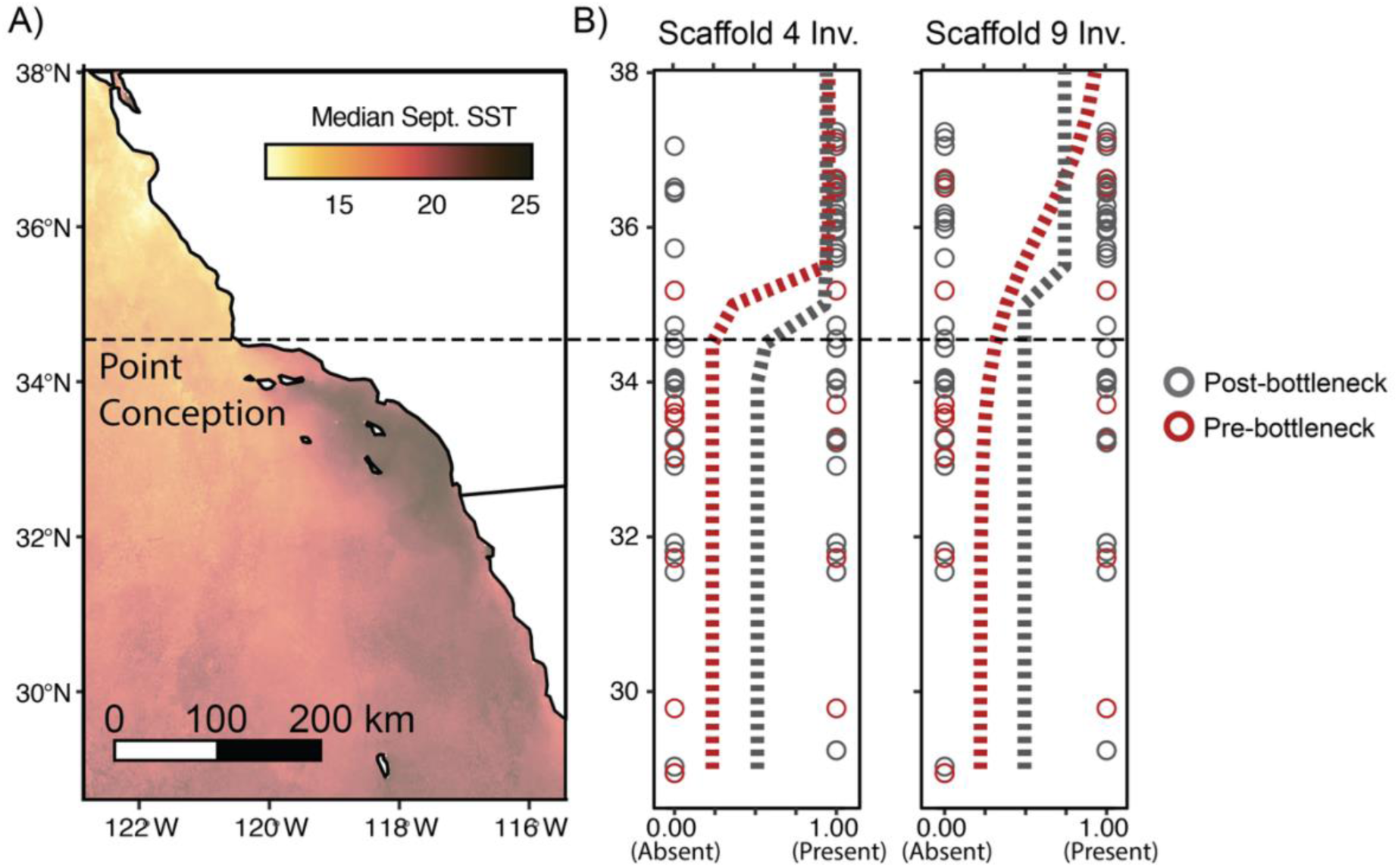
Inversion frequency changes after the Withering Syndrome bottleneck. **A)** Map of the California and Baja California coasts annotated with sea surface temperature and the Pt. Conception biogeographic boundary. **B)** Sigmoid clines fitted to inversion frequencies pre- and post-bottleneck. Each circle represents an individual and whether it has one or more copies of the collinear allele (0) or inversion allele (1).

The two inversion loci are in linkage disequilibrium with each other south of Pt. Conception. An initial analysis showed a significant correlation between having a copy of the scaffold 4 inversion (genotype = *A_*) and a copy of the scaffold 9 inversion (genotype = *B_*) south of Pt. Conception (r^2^ = 0.185; p = 1.66e-05). This correlation was absent at northern sites (r^2^ = 0.015; p = 0.22). Explicit tests of two-locus Hardy-Weinberg Equilibrium (HWE) show that, in post-bottleneck samples only, there is a significant difference between observed and expected inversion genotypes south of Pt, Conception (*X^2^*test, df = 6, p = 7.79E-04; Table S2). Surprisingly, this deviation appears to be driven by an excess of individuals without either inversion copy (genotype = *aabb*). We observe no appreciable deviation from HWE for any other combinations of time point and geography (p > 0.048). Together, these data suggest that, at least in the present day, epistatic interactions between the scaffold 4 and scaffold 9 inversion loci may be influencing their distributions.

## Discussion

Mounting evidence shows that time lags are common and can persist for long periods following a demographic bottleneck ^28^. For black abalone, we see that the nearly forty years following the Withering Syndrome bottleneck has been insufficient to change genetic diversity, inbreeding, or structure in a meaningful way. We hypothesize that, regardless of the original bottleneck intensity or any ongoing recovery, the ‘time-lag’ effect means that it is too early to see diversity loss, inbreeding depression, or population fragmentation take place. Fortunately, the perspective afforded by ancient DNA also suggests that black abalone may avoid future genomic erosion and that locally adaptive variation is being maintained.

### Shells as a genomics lens into the past

Our work represents the first population genomic dataset from mollusk shells and demonstrates that it’s possible to consistently and affordably generate whole genome data from this sample type. We attribute this success to the application of best practices in ancient DNA analysis. All work from shell powdering to library preparation was performed in a dedicated clean facility to reduce sources of contamination ^62,63^. Additionally, bleach treating the shell powder ^44,64^, performing extractions with silica spin columns optimized for small DNA fragment recovery ^50^, and using a single-stranded DNA (ssDNA) library preparation approach ^49^ produced complex libraries from low input DNA extracts. The combination of these elements allowed us to consistently generate multi-fold coverage of historic and ancient abalone shells, including a 34X genome from a 1500-year-old shell. Other recent analyses of shells, including experimentation with shell morphology and treatment methods ^44^ and the successful capture of nuclear loci from a 100,000 Ka mussel ^41^, confirm that shells are underutilized reservoirs of DNA. In future work, it would be useful to explore shell DNA preservation in relation to the specimen’s age, deposition environment, and shell mineral composition.

### Demography and selection in black abalone following Withering Syndrome

A time lag best explains why genetic diversity, inbreeding depression, and population structure have remained stable in black abalone following the Withering Syndrome bottleneck (Fig. 2 & Fig. 3)^14^. Even centuries-long delays to changes in heterozygosity, inbreeding, and genetic load are supported by theory ^12^ and have been documented in other threatened populations ^13^. Time lags are more likely to occur in black abalone because they mature late, have long life spans, and highly overlapping generations, all of which extend the influence of pre-bottleneck individuals over time ^28^. Because we were unable to incorporate these life traits into our simulations, the real time lag is probably underestimated (Fig. 4; see ‘Limitations’). Also contributing to time lag is the large pre-bottleneck Ne and range of black abalone.

Formerly large and diverse populations require more generations for heterozygosity to be impacted ^65^, and the estimated 99% reduction for black abalone would result in Ne greater than 3,000, well above thresholds suggested to minimize genetic drift ^66^ . Consistent with this, our 99% bottleneck simulations, though idealized, show that even at 0% recovery genomic erosion does not occur in the short term (Fig. 4A). For comparison, the timing and magnitude of myxoma virus spread in rabbits mirrors the WS bottleneck in black abalone, yet has also not significantly altered genetic diversity or population structure after roughly 70 generations ^32^. From this perspective, it’s unsurprising that we see stability across our whole time series (Fig. 2), despite the archaeological evidence for human impacts on black abalone over millennia of harvest67.

Our predictions of genomic erosion rest on accurate assessment of the WS bottleneck. The difference between a 99% bottleneck and a 99.9% bottleneck, the latter of which may better describe WS’s effect at select sites ^68^, has an impact on whether genomic erosion appears in the short term (Fig. 5). Under the more extreme scenario, only a 10% simulated recovery stabilizes genomic erosion by 100 generations post-bottleneck. A recovery rate of this magnitude has been recorded at some sites (Fig. S4), but is unlikely to be sustained for long. Related to this, our framework conflates Ne at the level of the species range with local (site-level) Ne ^66^, which may be lower (see ‘Limitations’). Conversely, the initial WS impact may have been overestimated if the disease was more harmful to higher intertidal abalone visible to surveyors^69^, or if gene flow from less impacted northern areas diminished the impact at southern sites ^3^. Increased migration, achieved through the indirect effects of population growth or individual translocation ^70^, may mitigate more serious impacts ^12^.

We also see evidence for natural selection on immune genes following the Withering Syndrome bottleneck (Fig. 5). Populations can adapt to intense selection pressures within just a handful of generations and leave genomic evidence of this process ^31,32,71^. Adaptation on these timescales is more likely to draw on standing genetic variation, and a selective sweep may involve multiple haplotypes carrying the adaptive allele. ^72^. If adaptation indeed occurred in response to Withering Syndrome, the high standing genetic diversity of black abalone suggests that selection would have acted on multiple haplotypes. Consistent with this scenario, we observe multiple distinct low-diversity alleles of the *SVEP1* outlier region, suggesting that multiple haplotypes carrying an adaptive mutation(s) may have undergone selection (Fig. 5D). In addition to this sweep signal, we observe large regions of high diversity and sequence divergence on scaffolds 7, 12, 14, and 19, which could indicate balancing selection ^54^. Short-term genomic signals of balancing selection are difficult to detect in contemporary samples ^19,54,73^, but the power of temporal samples to detect it remains unexplored except in experimental systems ^74^. Whether through balancing selection or other processes, the maintenance of diversity at immune loci is well documented in declining populations ^75,76^. The high diversity outliers we detect are enriched for genes involved in immunity across mollusks ^77,78^, in particular *DMBT1* (Fig. 5E) ^57^. What remains unclear is how diversity at loci like *DMBT1* has increased, as there has not been much time for *de novo* variation to arise. For *DMBT1* specifically, the tight clustering of pre-bottleneck individuals with those as old as 1500 BP (Fig. 5C vs. Fig. 5E), suggests that this result is not simply due to low sample size.

Other evidence for selection comes from the clinal shifts of the scaffold 4 and scaffold 9 chromosomal inversions (Fig. 6). Prior to this work, we had characterized the 31 Mb scaffold 4 inversion and its significant association with latitude, hypothesizing that it would be involved in local adaptation ^7^. With pre-bottleneck genomes, we now see that the scaffold 4 inversion has recently doubled in frequency at sites south of Pt. Conception - a common biogeographic barrier ^61^ - while remaining stable at northern sites (Fig. 6 & Fig. S7). Surprisingly, a second inversion locus on scaffold 9 shows the same pattern. These parallel shifts could result from interactions between inversion alleles, as supported by our observation of significant present-day LD at southern sites (Table S3). Inversions are common in wild populations ^79,80^ but instances of epistasis between inversions are few ^81,82^. Here it is tempting to speculate that epistatic interactions are shaping WS adaptation at the southern, more WS-afflicted sites ^3^. Alternatively, both inversions are responding independently to other selection pressures that vary with latitude and have changed on the same timescale ^80,83^. Regardless, the stark differences in inversion distributions across Pt. Conception and through time suggest that these loci may be important for local adaptation ^84^.

## Limitations of this study

Simulations of selected alleles and non-Wright-Fisher population dynamics (i.e. ^85^), would provide more confidence in our predictions for genomic erosion. Currently, incorporating these factors requires a forward simulation framework that records mutations and individuals over the course of simulation ^86^. This becomes prohibitively memory and time intensive when simulating the Ne of pre-bottleneck populations (∼3.0e5), and rescaling population parameters (e.g. Ne, μ) for computational speed can produce bias ^87^. Moreso, many aspects of black abalone biology remain uncertain because attempts to rear multiple generations in captivity have been unsuccessful. Larval duration and its effect on dispersal is uncertain ^6^, and dispersal ability will impact time lag ^28^, local Ne ^66^, and recovery. Finally, the functions of the putative targets of selection also remain uncertain. Functional genomic data that might validate candidate genes is limited for black abalone and its closest congeners, all of which are endangered. Sequence homology to experimental mollusk systems remains the best way to speculate on these selection targets.

### Implications for the conservation of black abalone

Future genomic erosion in black abalone is not guaranteed. If assessments of decline and recovery rates are accurate, even minimal growth may be sufficient to avoid the loss of genetic diversity, inbreeding depression, and loss of connectivity between sites. Knowing that genomic erosion is not a foregone conclusion can increase enthusiasm for conservation efforts ^85^ and better direct limited resources. Our genomic baselines show that black abalone lack genetic structure across much of their range, and that the diversity that existed prior to Withering Syndrome is still represented today. Therefore, captive breeding to preserve genetic variation may be premature, but translocations of reproductive adults from growing sites could be an effective and low-risk way of fostering recovery ^38^. While we do not know the fitness consequences of the inversions that segregate across Pt. Conception, a conservative strategy might restrict translocations to sites on the same side of this boundary. Whether considering inversions or other loci like *SVEP1*, these results demonstrate that data connecting genotype to phenotype to fitness are sorely needed for this species. Experiments to collect such data (e.g. transcriptome responses to heat stress) should be prioritized alongside other conservation efforts. This will require increased support and flexibility from state and federal regulatory agencies. Finally, while we find these results encouraging for the genomic future of black abalone, we would like to acknowledge this is only one component of conservation. Other components, for example the preservation of high-quality habitat, are essential to ensure that healthy populations will be around for future generations ^68^.

## Data Availability Statement

All raw sequence data will be deposited on NCBI’s SRA database upon formal publication of this work. Original code will be deposited at https://github.com/twooldridge/ShellGenomics.

## Competing Interests

The authors declare no competing interests.

## Funding

TBW was supported by a National Science Foundation Division of Ocean Sciences (NSF-OCE) Postdoctoral Fellowship (# 2307479). AAC was supported by the Universidad Autónoma de Baja California (UABC) 20th Internal Call. PR was supported by the National Marine Fisheries Service (NMFS).

## Acknowledgements

We thank Dr. Lindsey Groves from the LA County Museum of Natural History and Dr. Todd Braje from the University of Oregon for their help in providing abalone shells. We are also grateful for donations of shells from the Amah Mutsun Tribal Band with the help of Dr. Mike Grone. Computing support was also provided by the UC Santa Cruz Genomics Institute. Sequencing was performed at the UCSF CAT, supported by UCSF PBBR, RRP IMIA, and NIH 1S10OD028511-01 grants.

## Author Contributions

TBW conceptualized the project. TBW, JDK, SMF, WES, HCC, and TT performed shell DNA extraction and sequencing. TBW and JO performed all bioinformatics analyses. ZA executed the literature review of genes under selection. ALM guided the clinal inversion analyses. AAC sampled black abalone and provided DNA extractions. PR provided abalone population survey data. BAS provided funding and supervised the project. TBW wrote the manuscript with input and approval from all authors.

## Methods

### Sample collection

For pre-bottleneck samples, 44 black abalone shells were loaned from the Malacology Collection at the Natural History Museum of LA County, ranging in age from 1914 to 1979. 7 shells ranging from ∼800BC to ∼1880 BC were loaned by Dr. Todd Braje at the University of Oregon Museum of Natural and Cultural History, and 7 ranging from ∼570 BC to 1770 BC loaned from the Amah Mutsun Tribal Band. All shells were identified as black abalone based on morphology. For post-bottleneck samples, we combined 138 black abalone published in Wooldridge et al. (2024) with 16 samples from Baja California ^88^.

### Sequence data generation

#### Shells

All shell sequencing procedures were performed in a dedicated ancient DNA facility at UC Santa Cruz, following standard clean room criteria (Poinar and Cooper 2000). We first used a dremel to obtain a shell fragment from the anterior end of each specimen. For smaller or more fragmented shells we used whatever material was available. We then sampled approximately 50 mg of shell powder after pulverizing each shell using a Mixer Mill MM 400 (Retsch). We incubated shell powder at room temperature in a 0.5% bleach solution followed by three washes in 1 mL of molecular grade water to remove contaminants(Korlević et al. 2015; Boessenkool et al. 2017). Next, each powder sample was incubated overnight at 37 °C in digest buffer (1 mL: 0.45M EDTA, 0.25 mg/mL Proteinase K). Finally, DNA was isolated using the silica column-based method described in Rohland et al. 2018 using Binding Buffer D with a final elution of 35μL buffer EBT. All extracts were quantified using a Qubit 4 (Invitrogen) and the Qubit 1X dsDNA HS assay kit (Invitrogen).

Single-stranded library preparation was performed for all extracts following the protocol outlined in Kapp, Green, and Shapiro (2021), with modifications as described in Nguyen et al. (2023). Single-stranded libraries were indexed and amplified in 50 μL reactions containing 20 μL pre-amplified library, 25 μL AmpliTaq Gold 360 Master Mix, 2.5 µL of 20 μM i7 indexing primer, and 2.5 µL of 20 μM i5 indexing primer. Libraries were amplified in a Bio-Rad C1000 thermocycler using the following conditions: 95 °C for 10 m, followed by 11 to 19 cycles of 95 °C for 30 s, 60 °C for 30 s, and 72 °C for 60 s, followed by 72 °C for 7 m. Post-amplified libraries were purified using a 1.2X SPRI clean. Finally, libraries were visualized on an Agilent Fragment Analyzer. Libraries were pooled and sequenced on an Illumina NextSeq2000 at UC Santa Cruz (2 × 61bp) to assess endogenous DNA content. Libraries with sufficient endogenous DNA content and complexity were sent for deeper sequencing at UC San Francisco Center for Advanced Technology on an Illumina NovaSeq 25B (2 × 100bp).

#### Live samples

All modern samples in this study derive from collection efforts published in Wooldridge et al. (2024) and Delgadillo-Anguiano et al. (2025). DNA extracts from the Delgadillo-Anguiano study were provided by Dr. Alicia Abadia-Cardoso, and libraries were generated following the NEBNext Ultra II FS DNA Library Prep Kit for Illumina (NEB) using the recommended protocol with Y-Adapters in place of the NEBNext Adapters.. We incubated the samples for 5 to 6 minutes during enzymatic fragmentation, performed a single-sided 0.8X SPRI bead mixture prepared according to Rohland and Reich (2012). Libraries were amplified for 7 cycles using dual unique indexes. Libraries were eluted in 21 μL 0.1X TE and quantified using the Qubit dsDNA HS Assay (Invitrogen) and an Agilent Fragment Analyzer. All libraries were sequenced on an Illumina NextSeq 2000 before being sent for greater sequencing at the UC San Francisco Center for Advanced Technology on an Illumina NovaSeq 25B (2 × 150bp).

#### Alignment and variant calling

We adapter-trimmed and merged overlapping reads from all shell libraries using *fastp* with default parameters ^89^. Merging was necessary given the smaller average fragment size of DNA in the shell libraries. Modern abalone libraries were also processed by *fastp* with the read merging step omitted. Next, we aligned all shell and modern samples to the primary haplotype of the black abalone reference genome (GCF_022045235.1; ^90^ with *bwa mem* using default parameters ^91^. We elected to use the *bwa mem* approach for all sample types to reduce batch effects ^92^, as analyses with *mapDamage2* showed minimal damage and fragmentation in the vast majority of shells (Fig. 1). We marked and removed duplicates using *sentieon driver --algo LocusCollector --fun score_info* followed by *sentieon driver --algo Dedup --rmdup*. Finally, for all bam files we filtered for only primary alignments using *sambamba view -F "not (unmapped or secondary_alignment or supplementary)”*.

We next generated per-sample gVCF files using *sentieon driver –algo Haplotyper --emit_mode gvcf*. We then performed joint genotyping on this set of gvcfs using *sentieon driver –algo GVCFtyper --emit_mode ALL*, which produced invariant + variant sites across the genome. Finally, we filtered these variant sites using GATK VariantFiltration (Van der Auwera et al. 2013). We performed initial filtering on SNPs and INDELs independently, excluding SNPs with QUAL < 30.0, QD < 2.0, FS > 60.0, MQ < 40.0, MQRankSum < -12.5, ReadPosRankSum < -8.0 or SOR > 3.0 and excluding INDELs with QUAL < 30.0, QD < 2.0, FS > 200.0, ReadPosRankSum < -20.0, SOR > 10.0 ^93^. For invariant sites, we filtered based on site quality (“QUAL>30”). Finally, from this set of filtered variants we selected only biallelic SNPs at transversion sites. This transversion SNP dataset was used for all subsequent analyses unless otherwise specified.

#### Genomic masks

We generated three sets of genomic masks to remove regions that could bias downstream analyses.

First, we created a strict mappability mask with *GenMap* ^94^. We ran *GenMap* with -K 60 -E 2 to score regions based on mapping of 60mers with up to two mismatches. From this, we created a mask to exclude regions with scores less than 1. These excluded regions represent areas of the genome where only the most damaged and fragmented shell libraries would be susceptible to mismapping. Second, we created a mask to exclude two putative chromosomal inversions identified in Wooldridge et al. (2024). Third, we created masks based on depth. For this, we examined the distribution of site-level read depth in our variant + invariant site vcfs and excluded sites with less than 48 total reads (bottom 2.5% of sites) or more than 2884 reads (top 2.5% of sites).

#### Pseudohaploid genomes

As a complementary way to account for differential coverage across samples, we called pseudohaploid genotypes for all biallelic SNPs which passed quality and mappability filters by sampling a random read from each sample to represent genotypes at these positions. Pseudohaploid genotypes were called with SAMtools ^95^ v1.9 using *mpileup -B -q25 -Q30* and pileupCaller from sequenceTools v1.5.2 (https://github.com/stschiff/sequenceTools) with the *--randomHaploid* and *--singleStrandMode* options, which allowed for excluding genotypes calls potentially originating from ancient DNA damage. This enabled us to include transitions in downstream analyses. We created separate pseudohaploid genotype calls with major inversion regions included and excluded.

#### Population genetic analyses

All following population genetic analyses implement different strategies based on sequencing coverage and the extent of DNA damage. For clarity we have included a table outlining the sample sets and data types used for each analysis (Table S3).

### Genetic diversity and population structure

#### Heterozygosity

To analyze individual genetic diversity, we performed DNA-damage aware inference of heterozygosity in both full and 1X downsampled genomes using ROHAN ^52^. We first inferred DNA damage profiles for all shell samples using *bam2prof -minq 20 -both*. After confirming that the DNA damage profiles met those inferred by *MapDamage2*, we proceeded with heterozygosity inference using the command *rohan –rohmu 1e-4 –tstv 1.06 –size 500000 –chains 10000*, providing the deamination profiles with –deam5p and – deam3p. We then ran the same command for all modern samples, but without any deamination profiles. The choice of 1.06 for the transition-to-transversion ratio (--tstv) was informed by polymorphism in modern samples. For all commands, we also restricted our analyses to regions with good genome mappability (see Methods: Genome masks) with the *–map* flag.

#### PCA and inversion genotyping

We performed a genetic principal components analysis (PCA) on three data types for comparison: 1) genotype likelihoods, 2) pseudohaploid genomes, and 3) filtered variant calls. Given that all individuals clustered by inversion genotype in Wooldridge et al. (2024), and preliminary analyses with only high-coverage samples showed the same pattern, we had a strong *a priori* expectation that the inversion structure would be recapitulated in some form here.

However, an initial analysis of genotype likelihoods with *PCAngsd* ^96^ showed samples clustering strictly by whether the DNA came from shells or live individuals, regardless of coverage. The shell cluster also included the modern ‘control’ shell that we sequenced, leading us to believe that this structure might be artificial. To test this, we generated pseudohaploid genomes as is common with low coverage degraded DNA samples (see above), then pruned variants in these genomes to randomly sample 1 transversion SNP every 1 kb. We used these variants as input to PCA with *plink* ^97^ and *smartPCA* ^98^. Complementary to this, we obtained the variant calls themselves at these same pruned sites to serve as input to *plink* and *smartPCA* as well. Both *plink* and *smartPCA* returned similar structure on both the pseudohaploid genomes and the variant calls. This structure showed the effect of the inversion genotype in modern samples and grouped shell + modern individuals from outlier populations together (Fig. 3), leading us to believe that this PCA approach was accurately capturing population structure.

We then used our PCA analyses to assign inversion genotypes to each individual. Following Wooldridge et al. (2024), we generated PCAs based on variants from the scaffold 4 and scaffold 9 inversion loci. After confirming the three-cluster structure indicative of chromosomal inversions ^79^, we defined inversion heterozygotes (0/1) as individuals belonging to the central cluster (Fig. S8). Finally, we defined inversion homozygotes (1/1) as individuals belonging to the less diverse outer cluster, and reference homozygotes (0/0) as those belonging to the more diverse outer cluster.

#### Pi, FST & DXY

We generated diversity metrics from variant + invariant site VCFs with *pixy* ^99^, which estimates sequence diversity while accounting for the pitfalls in generating such estimates from heterogeneous data with high rates of missingness. We ran *pixy --stats pi, dxy, fst* on 50 kb x 25 kb sliding windows that were masked based on mappability and overall sequencing depth (see above). For comparison, we ran *pixy* for both the full sample set and high coverage subset, as well as on all polymorphisms and transversion polymorphisms only (Fig. S2). Given the similarity in pi (π) between high coverage pre- and post-bottleneck samples, we defined the statistic πpost-pre as the difference between post-bottleneck π and pre-bottleneck π. We used πpost-pre for further inference of differences in selection between the two time periods (see below).

### Genetic load

We estimated the load of deleterious mutations with *snpEff* ^100^ and *snpSift* ^101^. For this, we used the NCBI-generated annotation for the black abalone reference genome (GCA_022045235.1). We restricted this analysis to only high coverage (>15X) samples with confident genotype calls at transversion sites, allowing us to capture relative load between historic and modern samples. After scoring these variants with *snpEff*, we selected for loss-of-function mutations (filter "(exists LOF[*].PERC) & (LOF[*].PERC > 0.9)") and synonymous mutations (filter "ANN[0].EFFECT has ’synonymous_variant’") using *snpSift*. Finally, to obtain relative measures of load, we divided the loss-of-function variants by the number of synonymous variants for each sample.

### Gene content and enrichment analyses

We took shared genomic windows from our FST and πpost-pre analyses and intersected them with the *H. cracherodii* annotation from NCBI. We defined outliers as windows that were in the top 1% of FST and πpost-pre (e.g. balancing selection candidates) or the top 1% of FST and bottom 1% of πpost-pre (e.g. selective sweep candidates). We next added 5000 bp of to the start and end of each gene’s coordinates prior to this intersection in order to capture cis-regulatory regions under selection. We manually inspected each gene associated with an outlier window, and also used *Orthofinder* v.2.5.5 ^102^ to identify orthologous gene families with the more studied Eastern oyster *Crassostrea virginica* (GCF_002022765.2). For lack of a black abalone GO enrichment database, we manually tested for enrichment of scavenger receptor genes in balancing selection windows. To do this, we randomly selected 83 genes (the number in the 1% outlier windows) from the “gene” entries of the black abalone annotation and counted the number of times that the term “scavenger receptor” appeared. We repeated this 1,000 times to calculate significance.

#### Population genetic simulations

We used *msprime* for all coalescent simulations ^103^. We simulated the long-term demographic history for black abalone inferred by Wooldridge et al. (2025) and added a recent bottleneck and recovery (Fig. S3). We also generated a population split concurrent with the bottleneck to explore the effects of the bottleneck on isolation between subpopulations. Bottlenecks of 99%, and 99.9% intensity were applied, and these bottlenecks were followed by growth occurring at rates of -10%, -1%, -0.1%, -0.01%, 0%, 0.01%, 0.1%, 1%, and 10%. We set these growth rates to return to zero if Ne reached the pre-bottleneck state (∼3.0e5) or dropped to 100. All simulations were of 10 Mb chromosomes, and we generated 100 replicates of each parameter combination. We assumed a mutation rate of 8.60e-9 ^8^ and recombination rate of 1e-8.

For our short-term simulations (Fig. 4), we emitted diploid samples from this simulation at 10 generations prior to the bottleneck, the start of the bottleneck, and at intervals of 10 generations afterwards. From these samples we computed 1) heterozygosity, 2) fraction of the genome in runs of homozygosity (FROH), 3) FST between the pre- bottleneck baseline and post-bottleneck timepoints (Temporal FST), and 4) FST between post-bottleneck subpopulations (Subpopulation FST) (Fig. S3). Finally, we repeated the above procedure but calculated the same statistics at intervals of 100 generations post-bottleneck in order to investigate long-term effects (Fig. S5). All statistics were calculated with *tskit* and custom functions ^104–106^. Code for these steps is present in the scripts *expo_split_shortterm.py* and *expo_split_longterm.py*

#### Inversion clines and associations

Following inversion genotyping through PCA, we aimed to quantify spatial changes in the distributions of the scaffold 4 and scaffold 9 inversions over time. First, we applied a log transformation on the spatial data and a logistic transformation on inversion presence/absence following Westram et al. (2018)^107^. Then, we fitted stable, linear, and sigmoid clines, based on the formula encoded in the R packages HZAR^108^, to these data using a maximum likelihood search with *mle2* function from the R package *bbmle* ^109^. For the search parameter space, we limited the lower and upper bounds of the inversion frequency to -1e-5 and 1e5, the lower and upper bounds of the cline center to 28- and 37-degrees latitude, and the lower and upper bounds of the cline width at 0.1 and 10 degrees latitude. Following maximum likelihood estimation we determined the best fitting model of the three using Akaike’s Information Criterion (AIC) via the R stats function *AIC* (Table S1).

Complementary to this cline estimation, we tested for two-locus Hardy-Weinberg equilibrium and linkage disequilibrium between each inversion locus using a chi square test. Specifically, we calculated population-level allele frequencies for each inversion by timepoint and position relative to the Pt. Conception barrier (i.e. Pre-bottleneck and North). From these frequencies we calculated the expected genotype counts and derived the chi2 statistic using *sum((data$obs_counts - data$exp_counts)^2 / data$exp_counts)*. We then evaluated the significance of this statistic using the base R function *pchisq*(…,df = 6, lower.tail = F). All code for these analyses can be found in *cline_fitting.R*.

#### GBIF analysis

We aimed to quantify what proportion of all museum abalone specimens consisted of shells. To do so, we downloaded all Specimen records under TaxonKey “Mollusca” from GBIF on 21 October 2025. We removed entries with no listed preparation and filtered for entries with ‘Haliotis’ in the species field. We then summed the “IndividualCount” field for all specimens with a hit for the search term ‘*shell|dry|dried|concha|valve*’ in the ‘Preparations’ field.

## Notes

### Competing Interest Statement

The authors have declared no competing interest.

