## Supplementary Figures and Tables for "Ancient DNA from shells reveals delayed genomic erosion and rapid immune adaptation in the critically endangered black abalone"

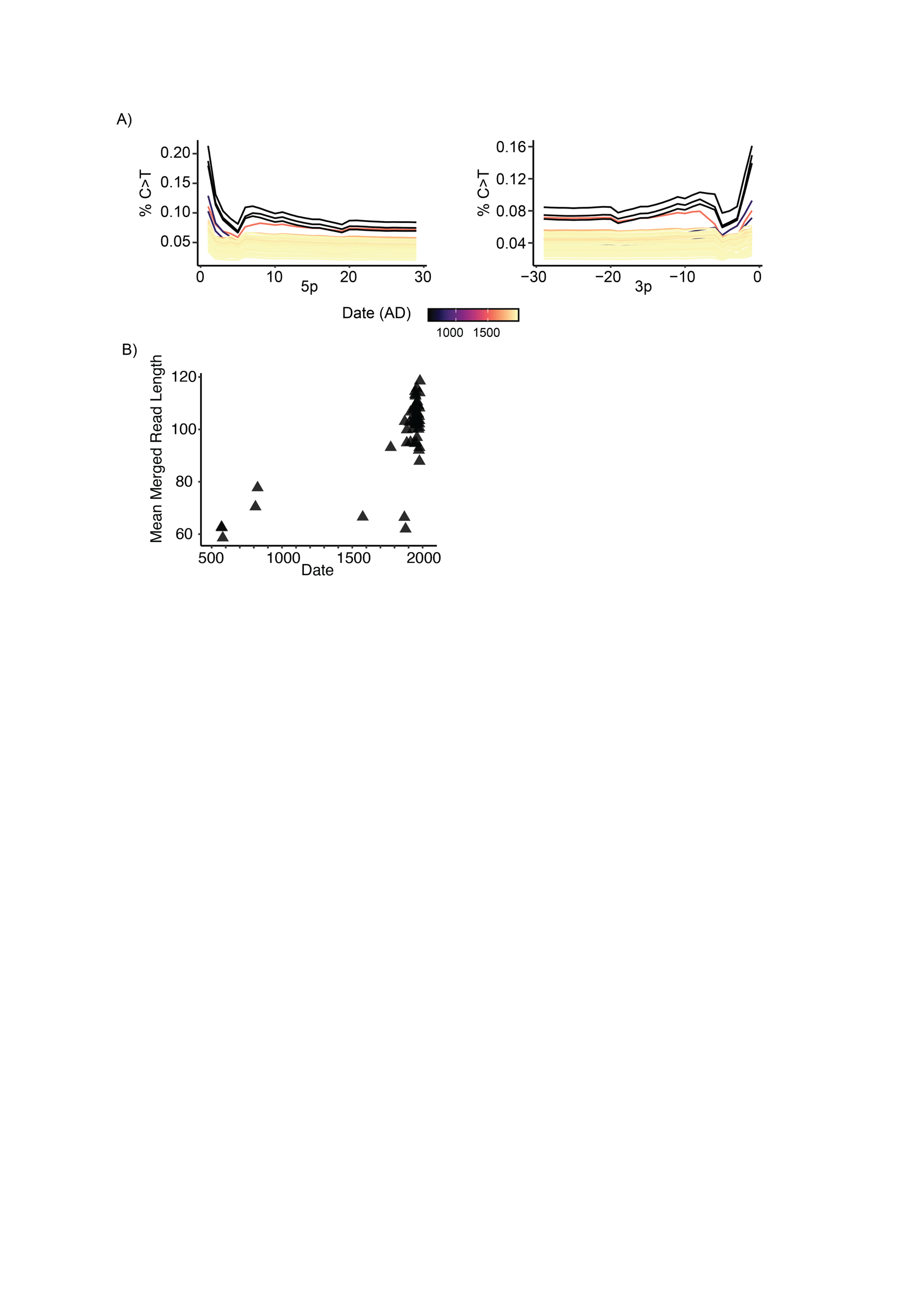
Supplementary Materials

**Figure S1. DNA Damage characteristics of shell genomic data. A)** Rates of cytosine deamination as characterized by *MapDamage*. **B)** Mean length of merged read fragments after processing with *fastp*.

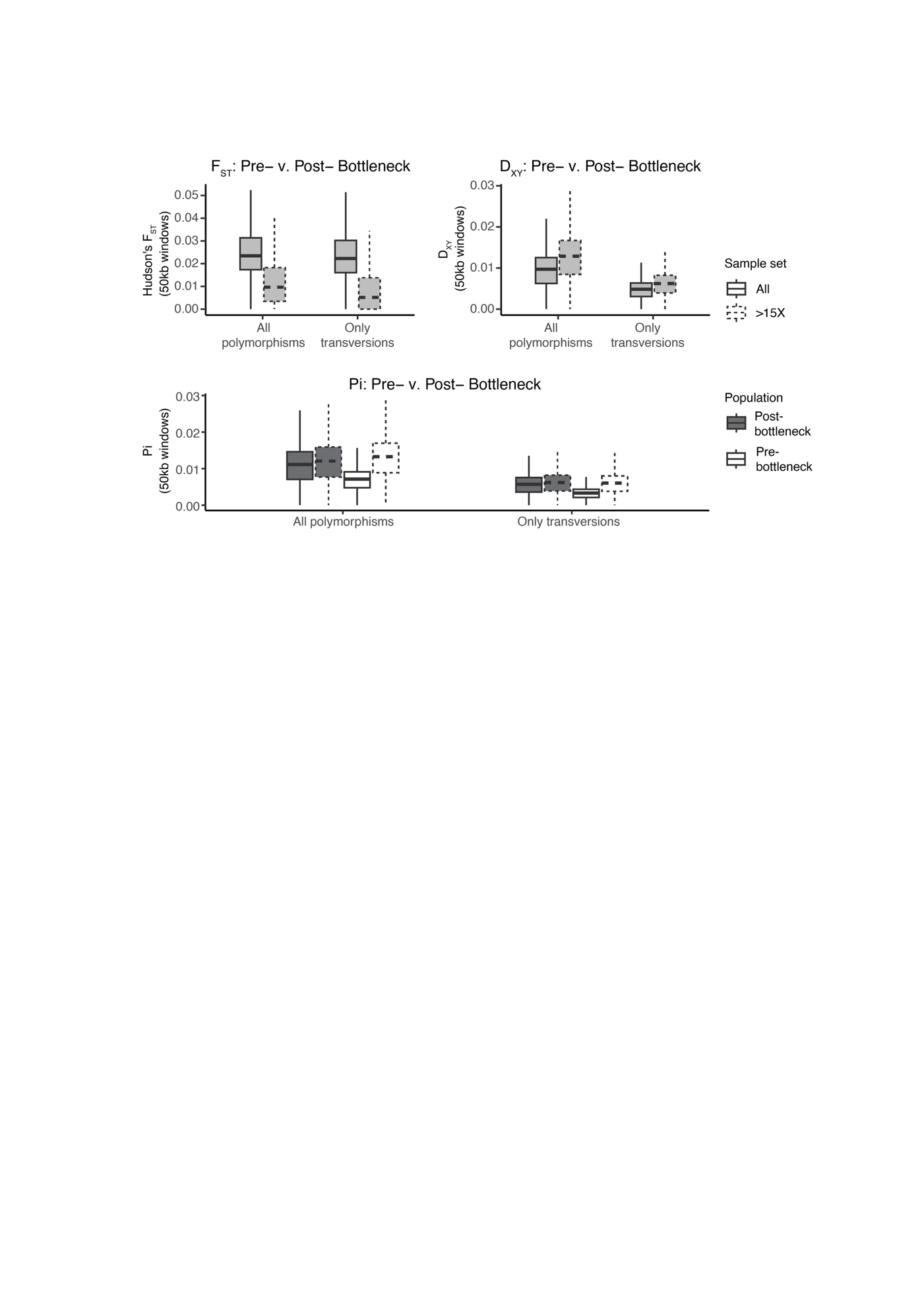

**Figure S2. Sequence diversity and divergence for pre- and post-bottleneck samples.** Statistics calculated on invariant + variant sites VCFs with *pixy* ^99^. Comparisons based on sequencing coverage and polymorphism type are shown to demonstrate the effect of reduced coverage, but not DNA damage, on relative D_XY_ and pi.

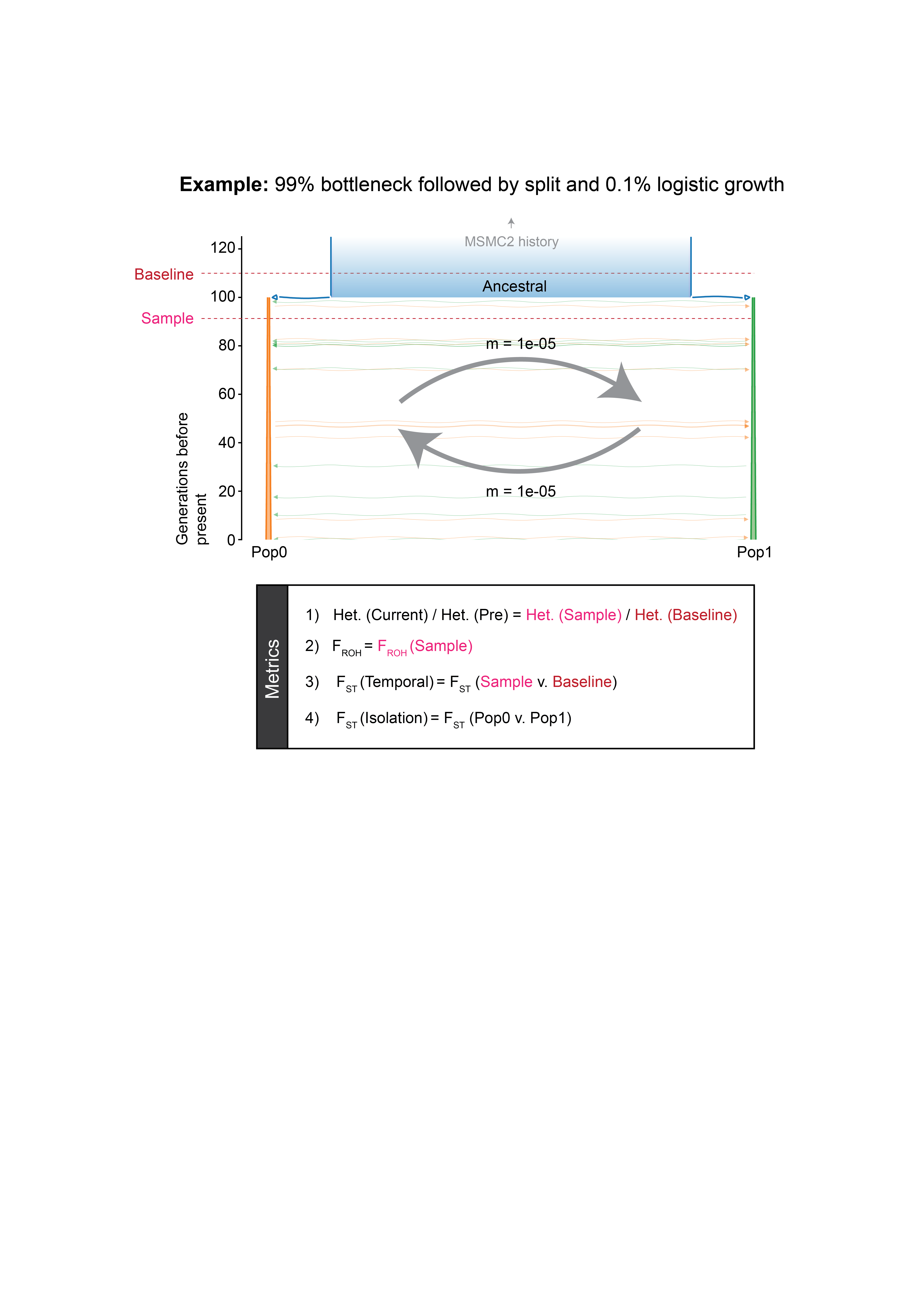

**Figure S3. Schematic of a simulated demographic model and population genetic metrics.** Only recent history is shown for clarity. The ‘Baseline’ timepoint always refers to 10 generations prior to the bottleneck. The ‘Sample’ timepoint here is shown at just 10 generations after the bottleneck but is also collected at regular intervals of 10 generations post-bottleneck up until the present. This interval changes to 100 generations in the long-term simulations. In the center, ‘m’ refers to the per-generation migration rate.

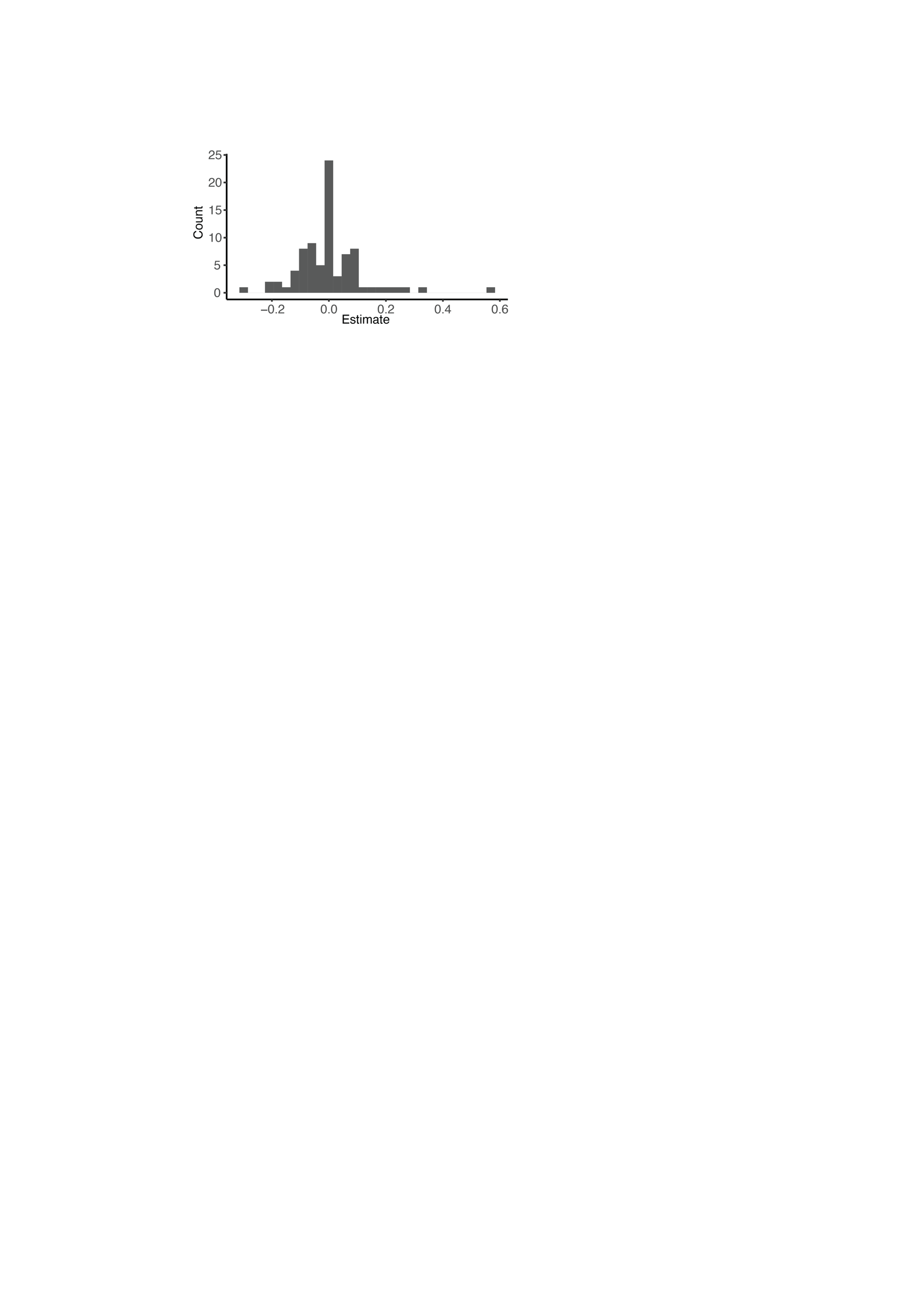

**Figure S4. Abalone population trends in California.** ‘Estimate’ refers to the 2018-2024 population trend at 82 sites. Data provided by Dr. Peter Raimondi.

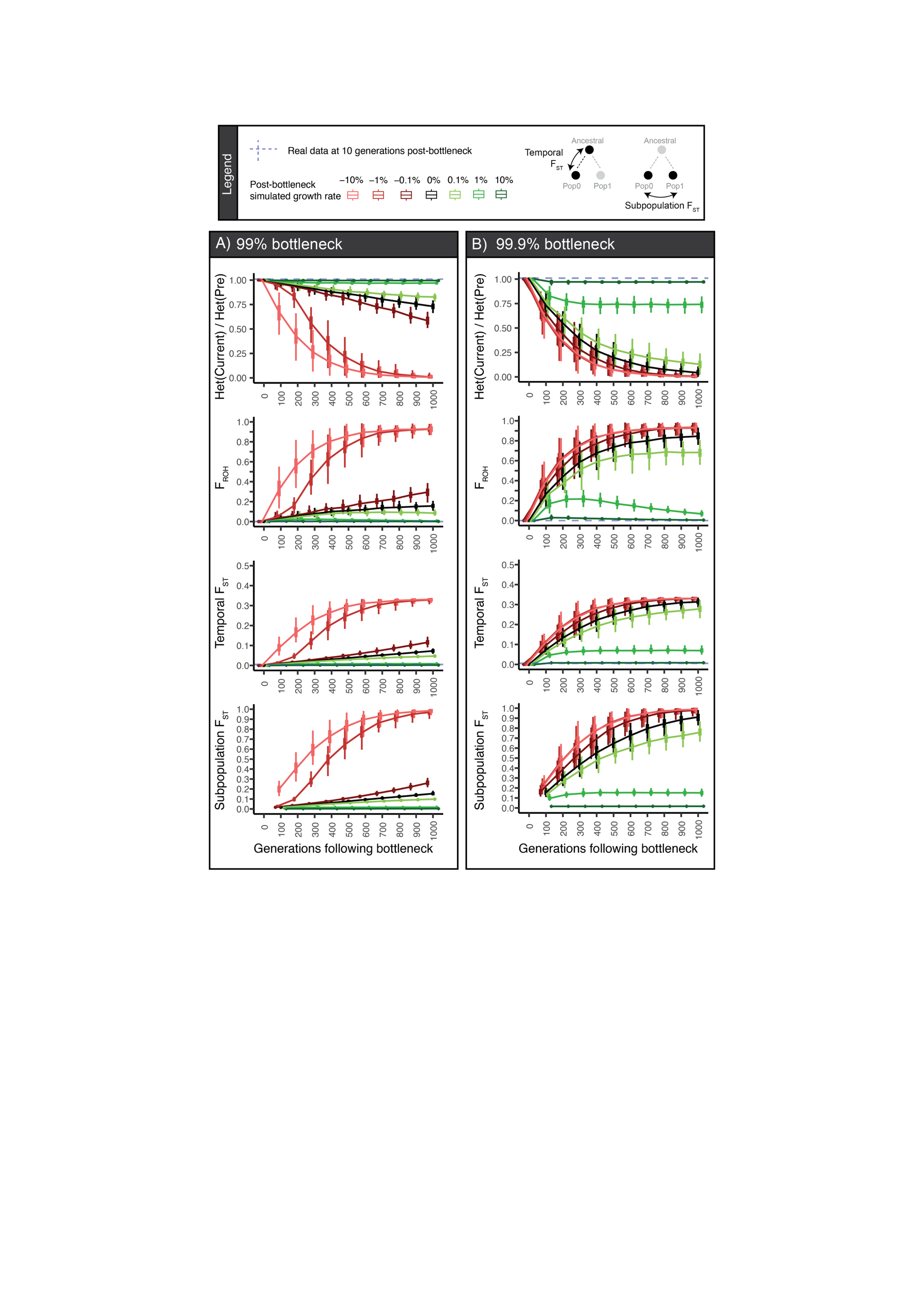

**Figure S5. Simulated effects of a bottleneck and recovery on inbreeding metrics.** These metrics include a ratio of heterozygosity at the current timepoint to 10 generations prior to the bottleneck (top row), fraction of the genome in runs of homozygosity (middle row), and Fst between the current timepoint and 10 generations prior to the bottleneck (bottom row). We simulated two bottleneck intensities, **A)** 99% of former size, and **B) 99.9%** of the former size. All growth and decline ceases if N _d_ returns to the pre-bottleneck Ne or drops to 100 individuals.

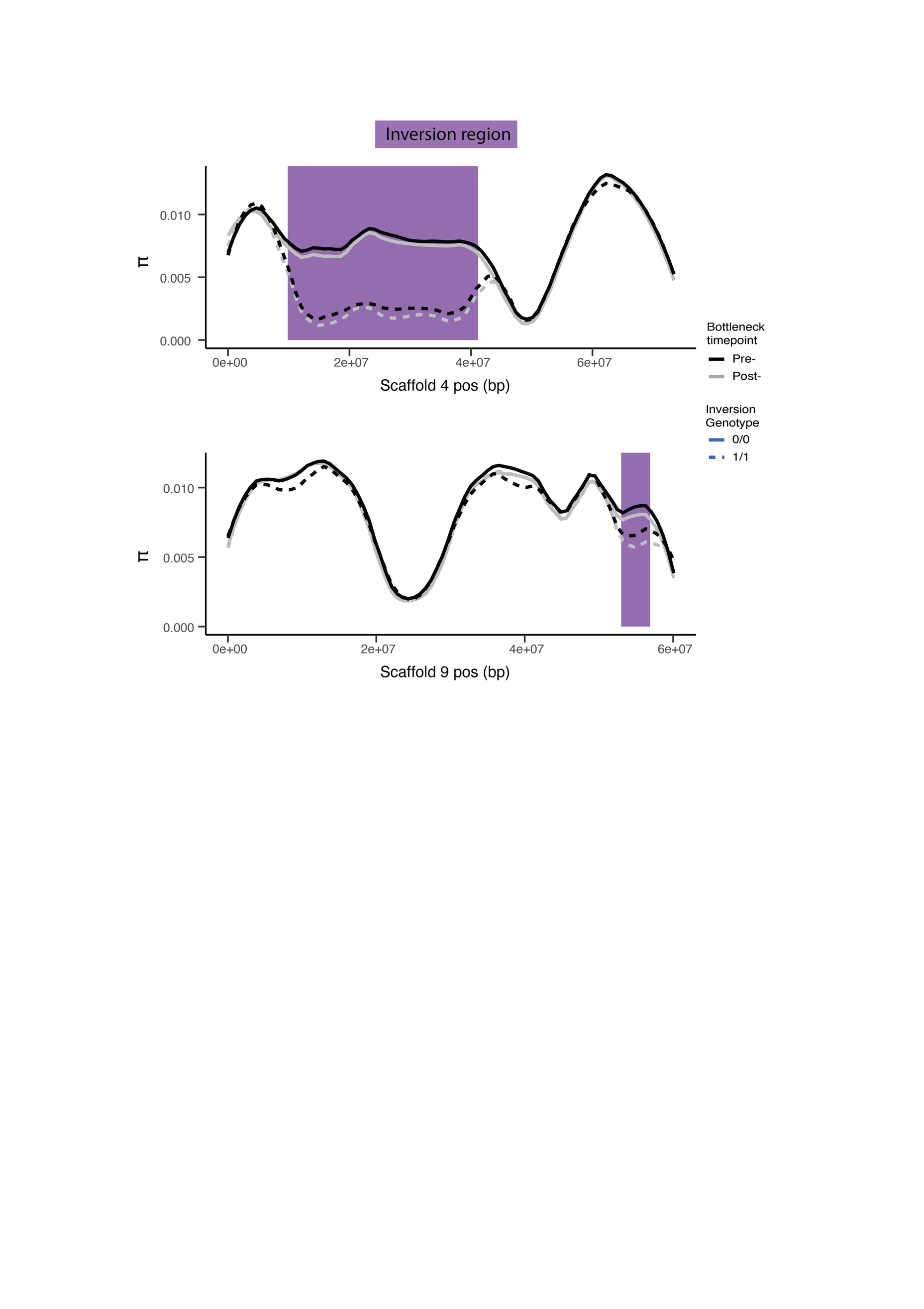

**Figure S6. Sequence diversity at the scaffold 4 and scaffold 9 inversions.** Loess-smoothed data of population-level pairwise diversity, with individuals broken into groups based on inversion genotype and sampling timepoint. No inversion heterozygotes are included.

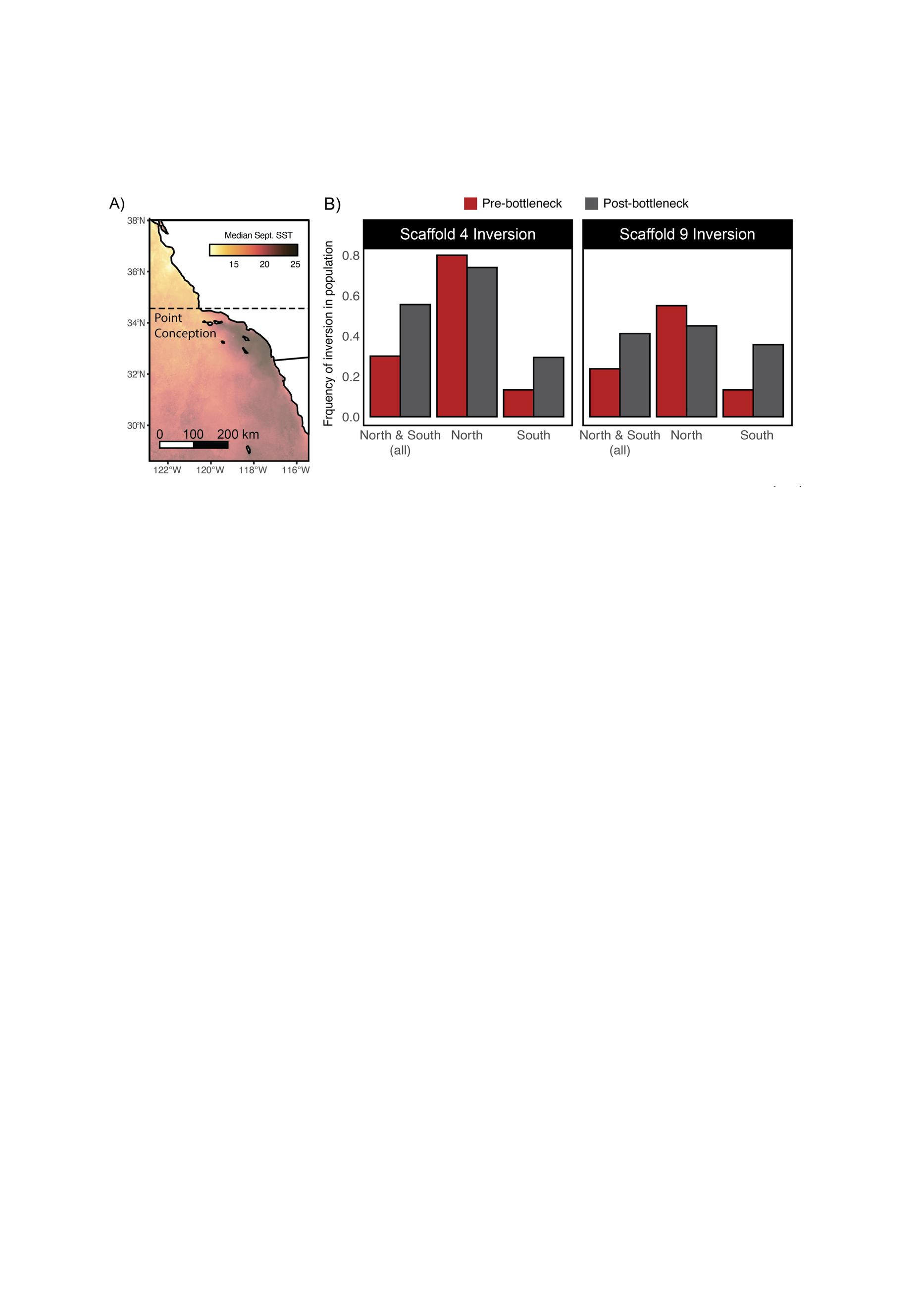

**Figure S7. Inversion frequency changes after the Withering Syndrome bottleneck. A)** Map of the California and Baja California coasts annotated with sea surface temperature and the Pt. Conception biogeographic boundary. **B)** Population level frequencies of both inversions relative to Point Conception. Bar heights indicate population-level frequencies of the derived (low-diversity) allele, presumed to be the inversion allele.

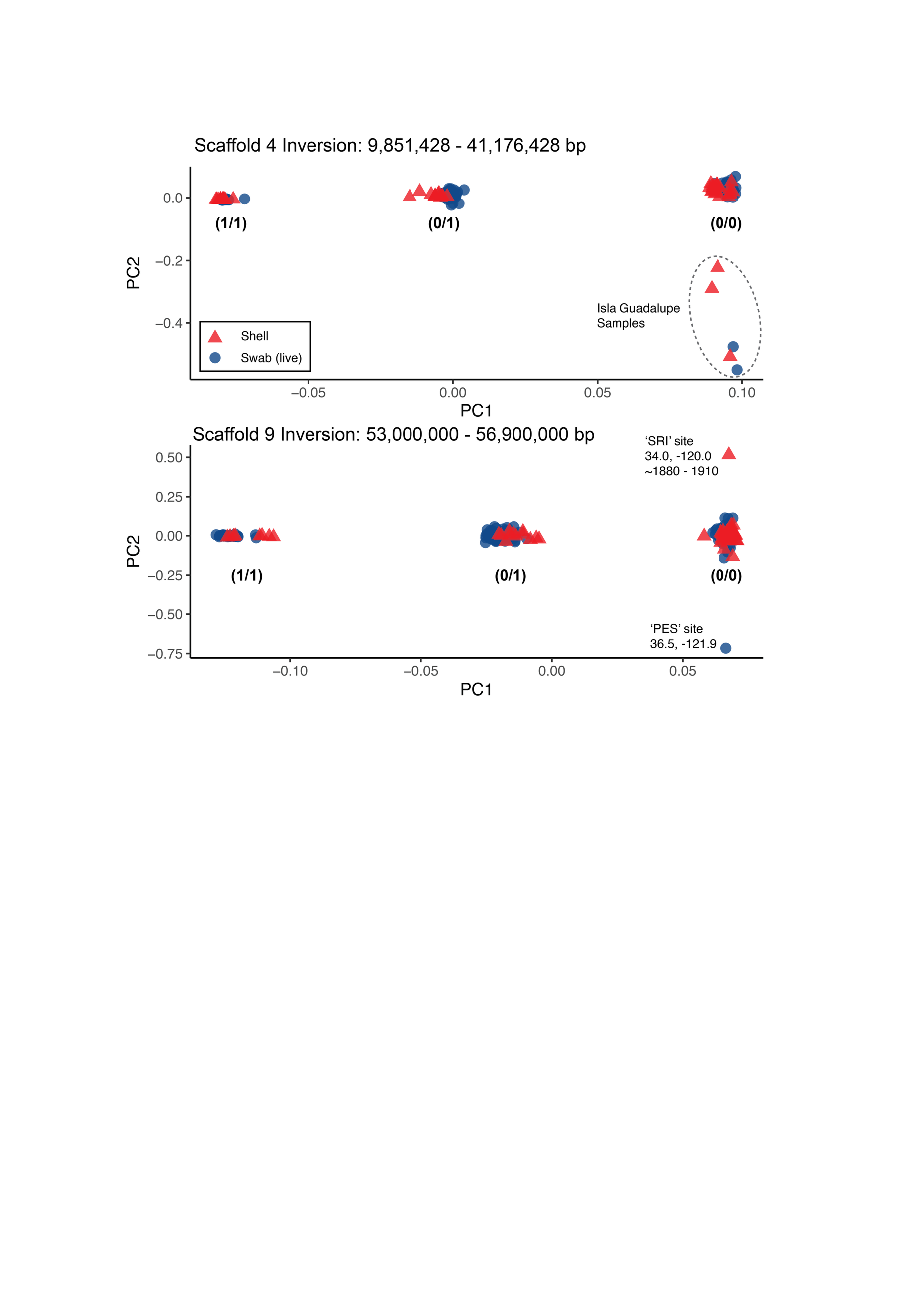

**Figure S8.** Genotypic PCAs at the scaffold 4 and scaffold 9 inversions. Each group is annotated with the assigned inversion genotype, 0 = reference allele and 1 = inversion allele. Outlier individuals are annotated.

**Table S1. Inversion cline fitting.** AIC Model comparison for inversion clines. “Inversion” = response variable indicating whether the inversion is present or absent in an individual’s genotype. “Time” = time period relative to bottleneck. “Model” = one of three clinal models. “AIC” = Aikake’s Information Criterion. “Df” = degrees of freedom.

| Inversion | Time | Model | AIC | df |
| --- | --- | --- | --- | --- |
| chr4inv_presence | pre | Stable | 55.84093 | 1 |
| chr4inv_presence | pre | Linear | 57.23144 | 2 |
| chr4inv_presence | pre | Sigmoid | 45.09568 | 4 |
| chr4inv_presence | post | Stable | 172.9499 | 1 |
| chr4inv_presence | post | Linear | 242.1008 | 2 |
| chr4inv_presence | post | Sigmoid | 142.4635 | 4 |
| chr9inv_presence | pre | Stable | 53.79573 | 1 |
| chr9inv_presence | pre | Linear | 57.07543 | 2 |
| chr9inv_presence | pre | Sigmoid | 52.90135 | 4 |
| chr9inv_presence | post | Stable | 203.1152 | 1 |
| chr9inv_presence | post | Linear | 249.1786 | 2 |
| chr9inv_presence | post | Sigmoid | 199.1223 | 4 |

**Table 2. Hardy-Weinberg Equilibrium analysis of inversions. “**Time” = time period relative to bottleneck. “Location” = location relative to Pt. Conception. “Genotype” = Two-locus inversion genotype, where AABB = homozygous for the scaffold 4 (A) and scaffold 9 (B) inversion alleles, and aabb = homozygous for the scaffold 4 (a) and scaffold 9 (b) colinear alleles. “Exp. Counts” = expected observations of each genotype assuming HWE and population allele frequencies. “Obs. Counts” = Observed numbers for each genotype. Below each Time x Location analysis are the A,a,B,b frequencies and the results of the Chi-square significance test.

| Time | Location | Genotype | Exp. Counts | Obs. Counts |
| --- | --- | --- | --- | --- |
| post | North | AABB | 9.953055 | 11 |
| post | North | AaBB | 7.03044 | 9 |
| post | North | aaBB | 1.241505 | 0 |
| post | North | AABb | 24.329691 | 25 |
| post | North | AaBb | 17.185519 | 10 |
| post | North | aaBb | 3.034791 | 6 |
| post | North | AAbb | 14.868144 | 14 |
| post | North | Aabb | 10.502262 | 14 |
| post | North | aabb | 1.854594 | 1 |
| A | 0.739 |  |  |  |
| a | 0.261 |  | Chi-square | 9.433 |
| B | 0.45 |  | degrees of freedom | 6 |
| b | 0.55 |  | p-value | 1.51E-01 |
| post | South | AABB | 0.6940195 | 2 |
| post | South | AaBB | 3.3331819 | 9 |
| post | South | aaBB | 4.0020857 | 0 |
| post | South | AABb | 2.5000252 | 1 |
| post | South | AaBb | 12.006924 | 11 |
| post | South | aaBb | 14.4164768 | 11 |
| post | South | AAbb | 2.2514233 | 2 |
| post | South | Aabb | 10.8129582 | 7 |
| post | South | aabb | 12.9829055 | 20 |
| A | 0.294 |  |  |  |
| a | 0.706 |  | Chi-square | 23.053 |
| B | 0.357 |  | degrees of freedom | 6 |
| b | 0.643 |  | p-value | 7.79E-04 |
| pre | North | AABB | 1.936 | 3 |
| pre | North | AaBB | 0.968 | 0 |
| pre | North | aaBB | 0.121 | 1 |
| pre | North | AABb | 3.168 | 3 |
| pre | North | AaBb | 1.584 | 0 |
| pre | North | aaBb | 0.198 | 0 |
| pre | North | AAbb | 1.296 | 1 |
| pre | North | Aabb | 0.648 | 2 |
| pre | North | aabb | 0.081 | 0 |
| A | 0.8 |  |  |  |
| a | 0.2 |  | Chi-square | 12.699 |
| B | 0.55 |  | degrees of freedom | 6 |
| b | 0.45 |  | p-value | 0.0481 |
| pre | South | AABB | 0.009387022 | 0 |
| pre | South | AaBB | 0.122384177 | 1 |
| pre | South | aaBB | 0.398898802 | 0 |
| pre | South | AABb | 0.122384177 | 0 |
| pre | South | AaBb | 1.595595207 | 3 |
| pre | South | aaBb | 5.200680617 | 3 |
| pre | South | AAbb | 0.398898802 | 1 |
| pre | South | Aabb | 5.200680617 | 2 |
| pre | South | aabb | 16.95109058 | 20 |
| A | 0.133 |  |  |  |
| a | 0.867 |  | Chi-square | 12.415 |
| B | 0.133 |  | degrees of freedom | 6 |
| b | 0.867 |  | p-value | 0.0533 |

**Table S3. Breakdown of population genetic methods and strategies.**

| Analysis | Data | Method | Notes |
| --- | --- | --- | --- |
| Heterozygosity and ROH | Bam files of all samples | ROHAN ^52^ | DNA damage and low-coverage aware method, useable for all samples |
| Genetic load | VCF of samples >15X, transversion sites only | SnpEff & SnpSift ^100,101^ | Requires confident variant calls, so high coverage only. Analysis included the most damaged (pre-1900) samples (Fig. S1), so transition were sites removed |
| PCA and inversion analyses | Pseudohaploid sequences of all samples, transversion sites only | Consensify ^110^ and PLINK ^111^ | Common pipeline for ancient/degraded DNA analysis. Allowed analysis of low-coverage and damaged data without relying on genotype likelihoods or low-quality variant calls. See Methods for details. |
| F_ST_, D_XY_, π summaries and sliding windows | VCF of samples >15X | pixy ^99^ | *Pixy* provides accurate calculation of π while accounting for missing data & invariant sites, assuming reasonable coverage. Calculations excluded most damaged (pre-1900) samples (Fig. S1), so no removal of transition sites. |
| Gene tree construction | VCF of samples >15X | IQ-TREE2 *^112^* | Maximum-likelihood inference requires confident variant calls, so only high coverage samples were used. |

**Extended Data Table 1.** All unique genes located in outlier windows from the selection analysis. Selection Type = Type of selection detected ; Gene symbol = Symobl of gene contained in 1% selection outliers, if applicable. Symbol inferred from protein description. “LOC” gene name retained if protein description was unclear. Protein Description = protein descriptions as per *Haliotis cracherodii* annotation on NCBI. Gene Function as per Mollusca Literature = Suggested gene function in shelled marine invertebrates (e.g. Abalone, clams, mussels, oysters) based on published studies, n/a means no relevant literature found. Primary Literature = Source publications for gene function annotations, n/a means no relevant literature found.
